# Seed encrusting with salicylic acid: a novel approach to improve establishment of grass species in ecological restoration

**DOI:** 10.1101/2020.10.27.356816

**Authors:** Simone Pedrini, Jason C. Stevens, Adam Cross, Kingsley W. Dixon

**Affiliations:** ARC Centre for Mine Site Restoration, School of molecular and life science, Curtin University, Kent Street, Bentley, 6102, Western Australia; Kings Park Science, Department of Biodiversity, Conservation and Attractions, 2 Kattidj Close, Kings Park, 6005, Western Australia; School of Biological Sciences, The University of Western Australia, 35 Stirling Highway, Crawley WA 6009, Western Australia; EcoHealth Network, 1330 Beacon St, Suite 355a, Brookline, MA 02446, United States; School of Molecular and Life Sciences, Curtin University, Kent Street, Bentley, WA 6102, Australia

**Keywords:** seed coating, seed enhancement, germination, emergence, grassland, demographic process

## Abstract

To achieve global ambitions in large scale ecological restoration, there is a need for approaches that improve the efficiency of seed-based restoration, particularly in overcoming the bottleneck in the transition from germination to seedling establishment. In this study we tested a novel seed-based application of the plant stress modulator compound, salicylic acid, as a means to reduce seedling losses in seed-to-seedling phase. First-time seed coating technology (encrusting) was developed as a precursor for optimising field sowing for three grass species commonly used in restoration programs, *Austrostipa scabra, Microlaena stipoides*, and *Rytidosperma geniculata*. Salicylic acid (SA, 0.1mM) was delivered to seeds via imbibition and seed encrusting with the effects tested on seed germination under controlled conditions (to test for resilience to drought), and in field conditions on seedling emergence, plant survival, and seedling growth. SA did not significantly impact germination under water stress in controlled laboratory condition and did not affect seedling emergence in the field. However, seedling survival and growth was improved in plants from SA treated seeds (imbibed and encrusted) under field conditions. When SA delivery mechanisms of imbibing and coating were compared, there was no significant difference in survival and growth, showing that seed coating has potential to deliver SA. Effect of intraspecific competition as a result of seedling density was also considered. Seedling survival over the dry summer season more than doubled when seed was sown at low density (40 plants/m^2^) compared to high density seeding (380 plants/m^2^). Overall, adjustment of seeding rate according to expected emergence combined with the use of salicylic acid is a cost-effective means for improving seed use efficiency in seed-based restoration.

## Introduction

Almost two-thirds of the world ecosystems are considered degraded or damaged with a lack of restorative effectiveness often unable to compensate for ecosystem loss [1]. Such degradation poses a serious risk to biodiversity, and impacts human communities that rely on ecosystem services for their sustenance and wellbeing [2,3]. Once degradation has occurred, restorative activities can be used to return the functionality, diversity, and structure of healthy, intact, and sustainable ecosystems [4,5]. Grasslands are among the most extensive terrestrial ecosystems in the world, covering over 52.5 million km^2^ [6], and provide fundamental ecosystem services such as sustaining food production (e.g., through rangeland pastoralism and dairy), carbon sequestration and storage, and erosion control [7]. However, almost half of the global grassland estate is considered degraded due to human activities and climate change [8] with important flow-on impacts for human societies whose livelihoods depend upon these grasslands.

In cases like of extreme disturbance, like post mining landscape, where spontaneous regeneration may not be feasible or effective, restorative interventions are required [9]. Native seeds of appropriate-local origin are commonly used to reintroduce missing species and to perform ecological restoration when the land has limited natural regenerative capacity [10,11]. However, abiotic factors such as nutrient-impoverishment, chemical and physically-hostile soil conditions [12] and low or unpredictable water availability [13], combined with biotic variables such as seed predation [14] and competition with exotic species, combine to limit the success of traditional seed-based grassland restoration.

Generally, less than 10% of sown native seeds become established plants, with significant bottlenecks detected at the seedling emergence phase [15], and in survival through the first summer drought [16]. Given the high cost and often highly limited availability of native seed [17], improving the efficiency in deployment to site is crucial if ecological restoration is to be delivered at the landscape scales expected [18] such as the UN Decade of Ecosystem Restoration. To address issues related to logistical constraints on seed delivery, and seedling establishment, the crop seed industry has developed technologies, such as seed coating, that could be adapted and applied to native seed [19].

Seed coating is the practice of covering seeds with external materials, sometimes including active ingredients conferring seeds protection and improved physiological performance [20]. Seed coating has been tested on native seeds in different restoration scenarios to overcome specific limitations such as water repellency [21], soil crusting [22], and seed predation [23]. However, despite promising results in seed coating improving seedling emergence, limited studies have so far attempted to improve native seed germination and seedling resistance to abiotic stresses [24].

Resistance to some abiotic stresses could be conferred by exposure of seeds to salicylic acid (SA). SA is a plant hormone, synthesised by many plant species [25]. It is involved in plant growth, developmental regulation [26], signalling [27], thermogenesis and mediating stress response either by providing resistance or triggering apoptosis [28]. Exogenous application of SA through watering, foliar spray, or seed imbibition has shown increased plant resistance and survival to a wide range of abiotic and biotic stresses [29]. SA efficacy in conferring stress resistance is a function of its concentration, with low concentrations failing to deliver resistance and higher concentrations decreasing resistance by activating cell death pathways [30,31]. The effect of SA on seed germination remains unclear; studies using seeds of crop species report improved germination for *Arabidopsis thaliana* under salinity stress [32] and for wheat (*Triticum aestivum*) under drought stress [33], while no effect was reported for maize (*Zea mays*) [34] or barley (*Hordeum vulgare*) [35]. Seed coating delivery of SA has shown some promising results when tested on tobacco seeds, improving germination and seedling growth under drought stress [36], and on corn, inducing resistance to chilling [37]. However, it has never been tested on native species for ecological restoration.

The goal of this study is to evaluate the effects of SA applied to seed on germination success, seedling emergence, survival and growth on three grass species native to southern temperate Australia, and to compare SA delivery methods via imbibition and coating.

The following hypotheses were tested: 1) coating or imbibition of seeds, without inclusion of SA, will not deleteriously impact seed germination success in laboratory trials or seedling emergence in the field, 2) SA will improve germination under conditions of water stress and enhance seed germination and seedling emergence in the field, and 3) plant survival and growth in the field will be improved for plants established from SA treated seeds at low and high intraspecific competition.

## Material and methods

### Species selection and seed processing

Three species of grasses native to temperate and Mediterranean regions of southern Australia were selected on the basis of their predominance in grassland revegetation and restoration activities and utility as pasture [38], including *Austrostipa scabra* (Lindl.) S.W.L. Jacobs & J.Everett, *Microlaena stipoides* (Labill.) R.Br. var. Griffin and *Rytidosperma geniculata* (J.M.Black) Connor & Edgar var. Oxley (all Poaceae). Seeds were sourced from a commercial provider (Native Seed Pty Ltd, Cheltenham, Victoria) in 2016. To reduce potential for viability loss seeds were stored in paper bags on open shelving in a controlled environment (15°C, and 15% relative humidity, RH) for one year prior to experimentation [39]. Seeds were moved to ambient condition (20–25°C and 40–50% RH) two weeks prior to experimentation to avoid potential seed damage during the cleaning and encrusting process [40].

Caryopses of each species were extracted from the husk to allow for more homogeneous encrusting and imbibition treatment. Removal of the palea and lemma was performed for each species using sulphuric acid digestion *sensu* Stevens *et al* 2015 [41], with complete immersion of the caryopsis in a 50% sulphuric acid solution (ACS reagent grade H_2_SO_4_, Sigma-Aldrich, St Louis, USA) for an optimal interval allowing for the weakening of floret structures without reducing germination potential. Immersion time for all three species was determined by Pedrini *et al* 2018 [42], and thus immersion intervals were 90 min for *A. scabra*, 60 min for *M. stipoides* and 20 min for *R. geniculata*. Acid immersion was followed by a neutralisation treatment in a 8.4 g L^-1^ sodium bicarbonate (NaHCO_3,_ Sigma-Aldrich, St Louis, USA) solution for 5 minutes, before rinsing under tap water for two minutes and drying in a Food Lab™ Electronic Dehydrator at 35° C (Sunbeam, Sydney, Australia). After drying, caryopsis extraction was achieved by gentle rubbing with a rubber mat and sequential sieving and zig-zag air flow separator (Selecta Machinefabriek BV, Enkhuizen, Netherlands).

### Seed treatments

After cleaning, caryopses (hereafter referred to as ‘seeds’) of each species were subjected to seed imbibition or coating treatments with or without salicylic acid application (Fig 1), resulting in four treatments (imbibed seeds without SA, imbibed seeds with SA, coated seeds without SA, coated with SA) plus an untreated control (uncoated, unimbibed seeds without SA). The coating treatment used in this experiment is defined encrusting, because the size and weight of the seed were increased but the shape of the seed remained evident [24].

**Fig 1.**
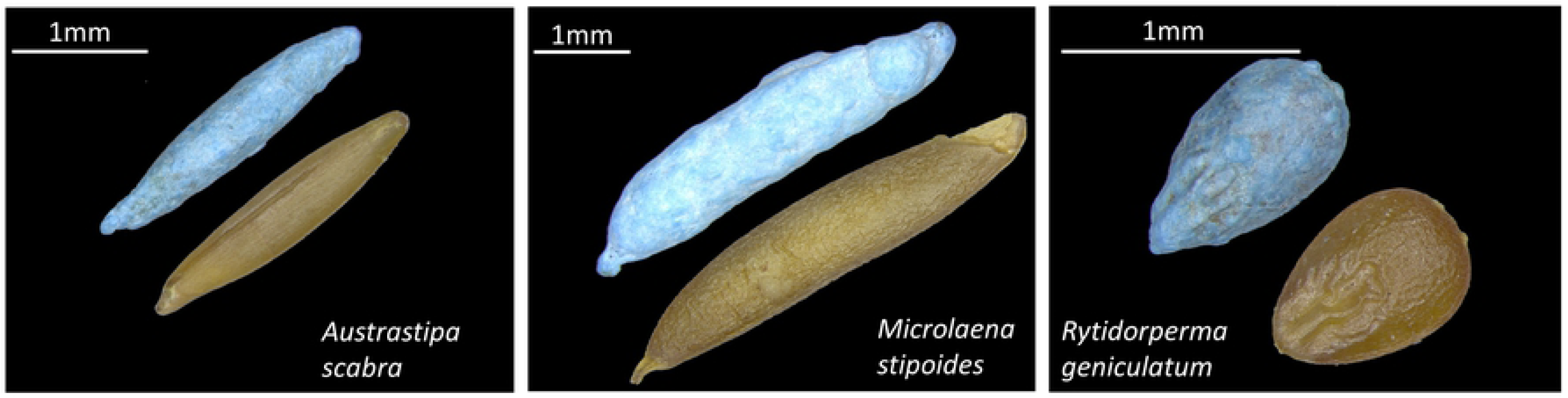
Seeds of the three grass species tested. In each image are presented the encrusted (blue) and untreated-imbibed seed. Scale bars indicate seed sizes.

SA was provided at a concentration of 0.1 mM, a concentration previously shown to be sufficient in confering stress resistance across various species and delivery methods [31,43,44]. SA solution was prepared by dissolving crystalline SA (Sigma Aldrich, St. Louis, USA) in deionized water for imbibition, and in a 2% Hydroxyethyl cellulose hydroxyethyl cellulose (cellosize QP 09-L, DOW chemicals) solution for encrusting (mixed with a magnetic stirrer for 30 minutes at 50°C). For imbibition treatments seeds were soaked in either SA solution or deionized water for 24 h at 20°C.

Seed encrusting was performed on a 15 cm RRC 150 Lab Coater (Centor Thai, Bangkok, Thailand), *sensu* Pedrini et al. (2018). Liquids were delivered through a compressed air-propelled 0.7 mm airbrush (Ozito tools, Australia). Talc was used as the filler material, dusted onto the seeds with a paint brush. Cleaned seeds (10 g) were placed inside the rotary coater, with rotor speed set at 300 RPM, and seeds were initially exposed to liquid spray until moist before powder was dusted onto the rotating seed mass. Wetting and dusting were repeated until 20 g of powder were used. A total of 15 ml of liquid were applied. Seeds were routinely checked to visually evaluate the even coverage of the coat, and to assess the formation of multiple seeds or dead balls (agglomerate of coating material not containing a seed). Following imbibition and encrusting treatments, seeds were placed on trays and dried for 3 hours in a in a Food Lab™ Electronic Dehydrator at 35° C (Sunbeam, Sydney, Australia).

### Laboratory test

Germination tests were performed in Petri dishes lined with two filter papers moistened with 14 ml water or Polyethylene Glycol (PEG) solution, placed in sealed plastic bags to reduce desiccation. 2 ml of water or PEG solution was added weekly.

In order to test whether SA improved germination success under water-limited conditions PEG 8000 (Sigma-Aldrich, St Louis, USA) diluted in deionised water at 24.72, 30.78, and 35.90 g/l was used to obtain solutions of −0.6, −0.9, and −1.2 MPa water potential at 20° C. This value resembles the range of water availability recorded in the field during the winter months. Germination tests were performed on four replicates of 25 seeds for each of the five seed treatments. Petri dishes were placed in a Biosyn incubator 6000 OP (Contherm, Korokoro, New Zealand) at 20°C with a 12 h photoperiod.

Germination was scored daily for the first five days and then at 7, 10 and 15 days respectively. On the 21st day, final germination was scored and remaining seed examined via cut test to assess viability. Non-viable seeds were excluded from the total.

### Field trials

Field trials were performed at a site east of the town of Waroona in Western Australia (32° 74’ 27” S, 116° 00’ 36” E, 201 m above sea level). The site falls within the native range of all three tested species and offers climatic conditions similar to those of mining operations active in the area likely to require these species in seed-based rehabilitation following mine closure. The field trial area was enclosed by a fence to avoid grazing from native marsupials and rabbits. Three experiments were performed in the field site: 1) seed germination in recoverable porous bags, 2) seedling emergence and survival in precision planted lines, and 3) plant survival and growth in plots. The five treatments previously described were tested in each experiment. For germination experiments in bags and lines, each treatment had four replicates. All experiments were arranged on a randomised complete block design of four blocks for 15 treatments (5 treatments * 3 species). For inline and plot experiments, seed were sown at depths of 0.2 − 0.5 cm, achieved by broadcasting dry soil on top of freshly sown lines and plots. All experiments were established at the commencement of the wet season in May 2017.

### Germination bags experiment

Field seed germination was tested by placing 50 seeds in 5 cm^2^ sealed mesh bags, over a 2 m^2^ area, and buried on site at 1cm depth. The bags were collected three weeks after sowing and germination recorded for those seeds as indicated by a protruding radicle.

### Line experiment – high competition

Seedling emergence was tested by sowing 100 seeds along a meter-long line, 5 cm wide. Seedling emergence was scored after 1, 2, 3, 4 6, 8 and 10 weeks. All emerged seedlings were left to then grow to maturity and resulted in high intraspecific competition. Plant survival was recorded 45 weeks after sowing.

### Plot experiment – low competition

To evaluate plant survival and growth under low intraspecific competition, 100 seeds were manually broadcasted on a 0.5 × 0.5 m^2^ plot. A month after sowing, the plots were thinned to 10 seedlings randomly selected, with at least 5 cm between seedlings, to limit potential competition resulting in a density of 40 plant/m^2^. The selected seedlings were marked with a pin to avoid confusion with other seedlings that could have emerged at a later stage. 45 weeks after sowing the surviving plants were counted, harvested and their height, wet weight and dry weight recorded.

Soil temperature and volumetric moisture content (m^3^/m^3^) were recorded for the duration of the germination and emergence experiment (10 weeks) with HOBO Micro Station Data Loggers (Onset Computer Corporation, Bourne, MA, USA). The probes were buried at 1 cm. For the 35 weeks following the end of the emergence experiment (July 2017 – March 2018), minimum and maximum temperature and precipitation data were obtained from the Dwellingup weather station, 10 km from the site [45] (**Fig 2**).

**Fig 2.**
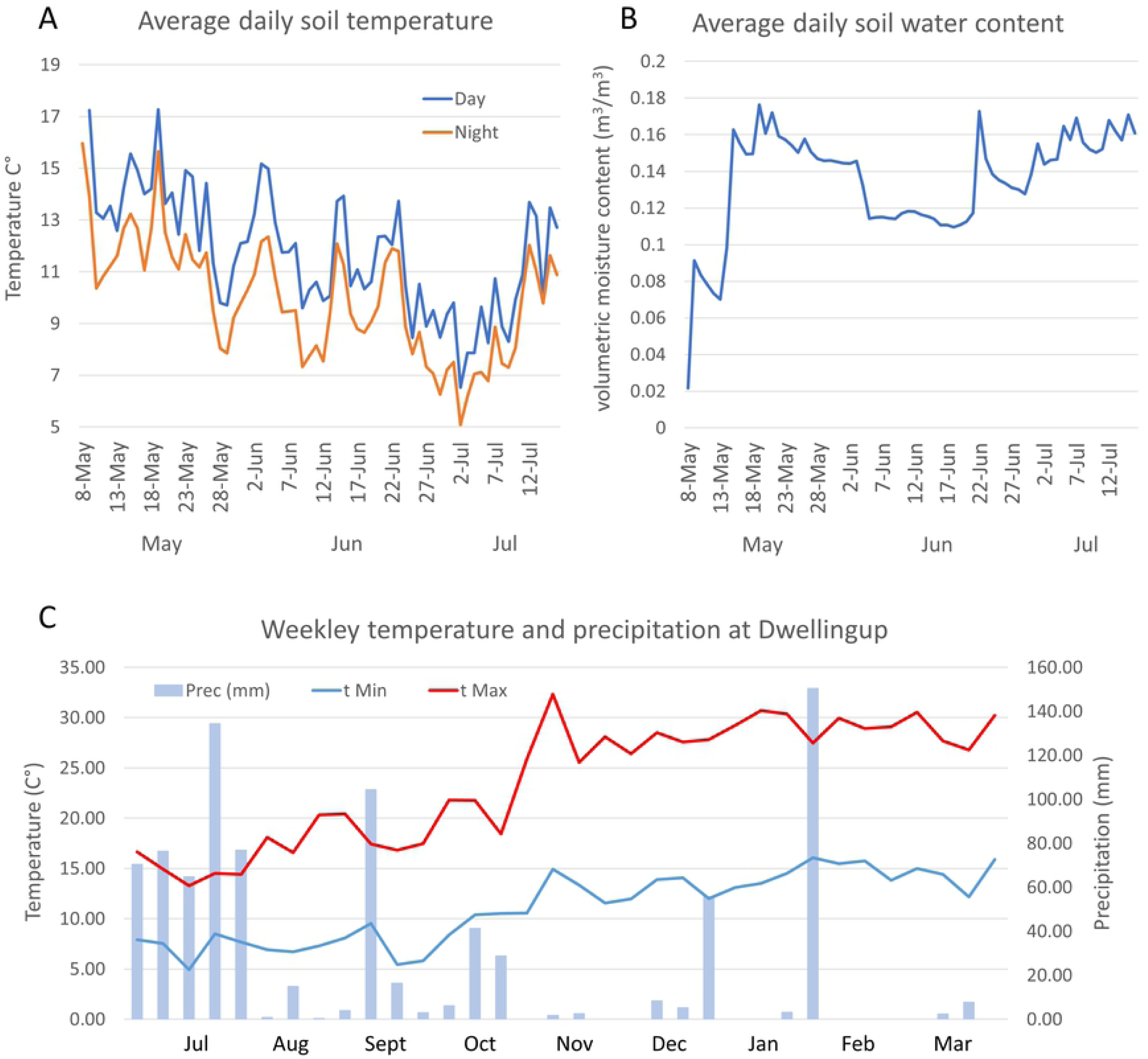
Climate condition at the field site. (A) the daily average for day (orange) and night (blue) temperature (B) volumetric water content in the soil at 1 cm depth for the first 10 weeks of the experiment, when germination and emergence were recorded. (C) Weekly maximum (tMax) and minimum (tMin) temperature, and total precipitation (Prec (mm)) for the period between the end of the emergence experiment and the recording of plant survival (July 2017 – March 2018) at a nearby meteorological station.

### Statistical analysis

To assess laboratory germination and seedling emergence in the field, non-linear regression models were fitted with the function “drm” of the “DRC” package [13,46,47]. A three parameter log-logistic model was used:

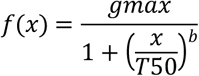

The parameters are: (b) slope curvature, (gmax) final germination and (T50) germination speed, intended as time (days/weeks) required to reach half of the final germination or emergence.

Parameter comparison on final germination and germination speed were then performed to assess differences among treatment (significance p <0.05).

To test the hypothesis of treatment and compound effect on germination in the field (in buried bags) and plant survival, an exact binomial test on the probability of success in a Bernoulli trial, between each treatment, was performed (confidence level = 0.95).

Plant height and biomass data were fitted in a Linear Mixed-Effects Model using the “lmer” function in the lme4 package for R [48], using compounds; untreated control (ctrl) vs treated without SA (NO) vs treated with SA (SA), and treatment; untreated control (ctrl), imbibed (Imb) and Encrusted (Encr) as fixed variables and the replicates (plots) as a random variable.

ANOVA (Type II Wald chi square tests) was employed to detect significant treatment effects. If such significance was detected a pairwise t-test was performed to compare the levels within the treatment. All data analysis was performed in the R statistical environment [49].

## Results

In the first two sections are reported the results of seed germination under laboratory conditions and seed germination/emergence in the field experiment, with the third section covers plant survival and growth data, collected at the field site.

### Encrusting and imbibition treatment

Encrusting treatment (Encr) had higher or similar germination than the control (Ctrl), whilst imbibition treatment (Imb) at times resulted in lower germination. Final germination of *A. scabra* treated seed, tested in lab conditions, was not significantly different from the untreated control, and only slightly but significantly (P < 0.001) increased in germination speed (T50) of 0.5 days, for both imbibed and encrusted seed. When tested in field conditions, the encrusted seed had lower final emergence than the control (Ctrl: 52 ± 1.6%, Encr: 45 ± 2.4%, P < 0.001) while imbibed seeds showed no significant difference (**Fig 3**).

**Fig 3.**
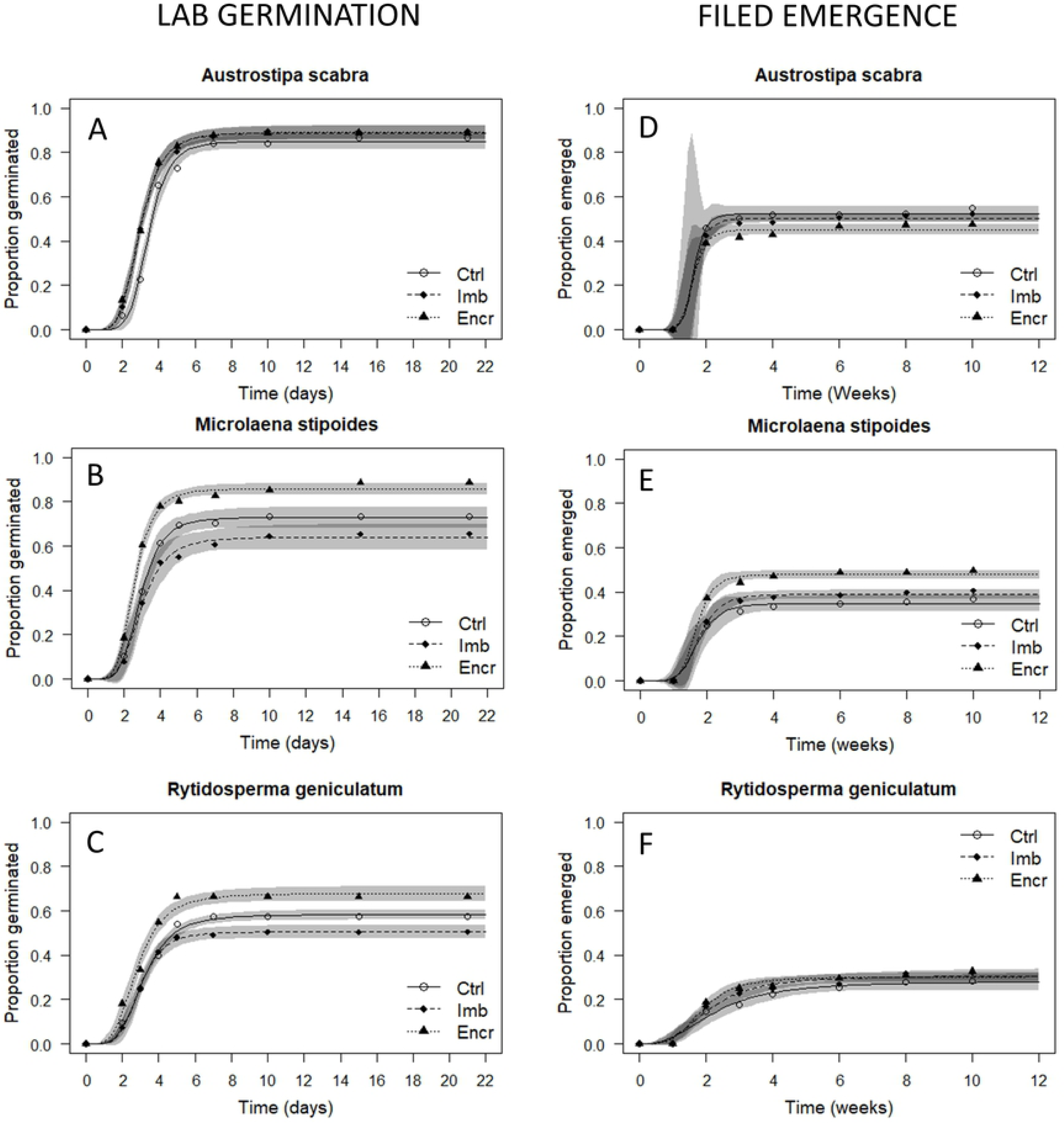
Seed treatment germination and emergence curves. Cumulative germination/emergence percentage curves of the three different seed treatment tested: untreated (ctrl), encrusted (Encr), and imbibed (Imb) across the three species tested. The lines represent the cumulative germination curve over time. Data points are the germination recorded on a specific day/week and the shaded areas represent the 95% confidence intervals. A, B and C germination experiments were in controlled laboratory condition. D, E and F seedling emergence in the field trial.

Under laboratory conditions encrusted *M. stipoides* seeds (86 ± 2.1%) germination was higher than in the control (73 ± 2.2%, P < 0.001), but 8.9% lower for imbibed seed (P < 0.05). Similarly, final emergence in the field was higher for encrusted seed (Encr: 48 ± 1.0%, Ctrl: 35 ± 1.0%) with imbibition increasing emergence by 4% (P < 0.05).

As with *M. stipoides*, germination of *R. geniculatum* was significantly higher for encrusted seeds (68 ± 1.5%) with the lowest for imbibed seeds (51 ± 1.4%), (Ctrl: 58 ± 1.5%). However, there was no difference in seeding emergence in response to seed treatment under field conditions.

### Salicylic acid effects on germination with low water availability and field emergence

To assess the effect of SA, seeds that were provided SA (via imbibition and encrusting) were compared to seeds that received the treatments without SA (NO). If a significant difference was detected, SA delivery methods of encrusting (ES) and imbibing (IS) were then compared. The high variability in the results suggested that SA has limited effects on promoting germination and emergence.

Final germination at optimal water potentials in *A. scabra* was significantly (P < 0.05) reduced by 4.3% with SA treatment (**Error! Reference source not found**.). At reduced water availability of −0.6, − 0.9, and −1.2 MPa SA treatments generally showed a slight but non-significant improvement in final germination. When tested in the field, SA treatments did not affect germination but reduced final emergence (NO: 51 ± 1.1%, SA: 44 ± 1.1%, P < 0.001) with SA encrusted seed emerging 5.6% lower than SA imbibed seeds.

Similarly, *M. stipoides* germination at optimal conditions was reduced in SA treated seed by 7.9% (P<0.05). SA delivered through encrusting resulted in better germination (77 ± 2.1%) than SA imbibed seed (57 ± 2.2%). Under limiting water potentials of −0.6 MPa, germination for SA treated seed was improved from 77% ± 1.9% to 86 ± 1.9%, and encrusting allowed for a 12.7% increase in germination compared to imbibing. However, at lower water potentials, SA treatment reduced final germination by 5.6% (P < 0.05) at −0.9 MPa and by 11.2% (P < 0.01) at −1.2MPa. In both situations encrusting allowed for better germination then imbibition. Field germination and emergence of *M. stipoides* were not significantly affected by SA treatment, but both treatments had higher emergence than the untreated control.

When final germination was tested on *R. geniculatum*, no significant difference between seed treated with and without SA was detected at optimal conditions and with reduced water availability. The only effect of SA was a delay in germination at 0.0MPa of 0.4 days. Field germination was no different for seed treated with and without SA, however both treatments had lower germination than the untreated control. Between seeds treated with and without SA, there was no difference in field emergence. However, seed treated without SA had significantly lower germination (p<0.05) than the untreated control. Emergence in SA treated seeds was slightly higher, but not significant.

The results of germination and emergence experiment are provided in the supplementary file S1_GerminationEmergenceAnalysisResults.pdf.

### Survival and plant growth in field site conditions

Plant survival was examined in situations where intraspecific competition was maintained high (line experiment) or reduced (plot experiment). In both scenarios, SA improved plant survival and growth. In the “line experiment” the survival of plants that emerged from untreated seed was 32.3% for *A. scabra*, 41.2% for *M. stipoides* and 42.6% for *R. geniculatum*. Plants emerging from SA treated seed, compared to seeds treated without SA, had a significantly (P>0.001) increased survival by 12.9% in *A. scabra*, 13.5% in *M. stipoides* and 11.8% in *R. geniculatum*. In *A. scabra*, SA delivered through encrusting improve survival by 9.8% (P>0.001) compared to SA delivered through imbibing. In *M. stipoides* and *R. geniculatum*, no difference was detected between SA delivery systems on plant survival.

In the plot experiment, the average survival of seedlings in the untreated control was of 82.5% for *A. scabra*, 82.5% for *M. stipoides* and 77.5% for *R. geniculatum*. In SA treated *M. stipoides* and *R. geniculatum*, compared treated without SA, survival was significantly improved (P<0.01), by 8.2% and 15% respectively and in *A. scabra*, survival was improved by 6.25%, but the difference was not significant. SA delivered through encrusting provided slightly better but non-significant survival. Both for *M. stipoides* and *R. geniculatum*, SA treatment improve survival by 17.5% and 10% respectively, compared to seed treated without SA (**Error! Reference source not found**.).

Plant growth was recorded in term of plant height and above ground dry biomass. In *A. scabra*, no significant difference was detected between SA and non-SA treatments in either measurement. For *M. stipoides*, plant height for SA treated seed was significantly improved (P<0.05) from 41 cm ± 1.7 cm (untreated control) and 43 cm ± 1.0cm (treated seed without SA), to 46 cm ± 1.0 cm. Dry above-ground biomass was also higher in SA treatment (3.4 g ± 0.22g) compared to untreated controls (2.2 g ± 0.25 g) and without SA (2.7 g ± 0.25 g) (both P<0.05). In *R. geniculatum*, there was no significant difference in height. Dry biomass for SA treatment (1.5 g ± 0.08g) was significantly higher (P>0.05) than treated without SA (1.2 g ± 0.10g), but not significant compared to the untreated control (1.3 g ± 0.09g). No significant difference between SA delivery through imbibing or encrusting, in terms of plant growth, was detected in the study species.

## Discussion

### Seed treatment effects on germination and emergence

Of the three species tested, only *A. scabra* showed no treatment (encrusting and imbibition) effect on germination and emergence as predicted. *M. stipoides* and *R. geniculatum* showed unexpected, significant differences between treated seeds (imbibed and encrusted) and the control. In the germination experiment, the two species behaved similarly, with encrusted seeds performing better than controls, while imbibition had negative effects on both final germination and germination speed. In this study, seeds were imbibed for 24 hours, following previously described methodology for SA delivery to seeds [33,50]. A potential explanation for the reduction in germination of imbibed seed could be anoxic stress due to extended submersion in water and in a water-saturated environment (petri dish). This problem has been reported in seed priming treatments that rely on seed imbibition to trigger pre-germinative metabolic mechanisms [51,52]. Oxygen availability could also explain why encrusted seed performed better than imbibed and untreated seed. During the encrusting process, seed contact with water was limited compared to imbibing. Moreover, the layer of encrusting material could also have acted as a buffer, reducing the water potential at the seed level and allowing for improved gas exchange. Furthermore, the emergence of imbibed seed was unaffected in the moist, but not water-saturated soil conditions. In seed priming treatments, water potential or water oxygenation are usually regulated [53] to avoid anoxic damage. The germination reduction detected in this study for imbibed seed could, therefore, be mitigated by decreasing imbibition time, reducing the water potential, or providing oxygenation to the solution.

### Salicylic acid effect on seed germination and emergence

Contrary to what was initially hypothesised, SA application did not clearly improve seed germination and emergence in the field and in controlled laboratory condition across a water availability gradient on the tested species, with the exception of *M. stipoides* at −0.6 MPa. *M. stipoides* seed treated with SA had significantly lower germination at 0.0, −0.9 and −1.2 MPa, suggesting that this species might be susceptible to the SA concentration tested. Germination response to exogenous SA application is concentration dependent, with inhibition detected at higher concentrations [35]. Reducing SA concentration for *M. stipoides*, could therefore potentially remove the germination impediments. When a difference in germination was detected for seed treated with SA, encrusted seed performed slightly better than imbibed seeds. However, this difference is most likely due to the process itself, as highlighted previously, other than the efficacy in delivering SA.

A significant drop in emergence by SA treated seed in *A. scabra* might suggest that the interaction of SA treatment with unidentified variables present in the soil at field site might have triggered a negative response, similar to what was observed in the controlled lab environment. Moreover, the detrimental effect of encrusting could have been determined by the combined effect of SA and the physical constraint of the coatings layer and soil to the emerging seedling. However, this effect was not detected in the other species.

### Survival and growth

In experimental plots where competition was reduced, plants from seed treated with SA resulted in increased height and biomass production in two out of the three species tests. SA also provided a significant improvement in plant survival in both scenarios with and without interspecific competition. Although response among species varied, with the least effects detected in *A. scabra*, the overall trend showed marked benefits in term of survival and plant grown from SA-treated seeds. The improved survival at this stage could be explained by the already described stress resistance proprieties of SA [44]. A potentially significant, yet unintended, result of this experiment is the great difference in plant survival between the low and high seedling density (line and plot experiment). According to the seedling emergence data, the seedling density in the line experiment was of 520 seedling/m^2^ in *A. scabra*, 430 seedling/m^2^ in *M. stipoides and* 280 seedling/m^2^ in *R. geniculatum*, whilst for the plot experiment seedling density was 40 seedling/m^2^ across all species. Based on personal observations, the plants with limited competition were generally more developed before summer than the ones in the lines. This would have allowed for the development of a broader and deeper root system with better access to water during the dry summer months ultimately resulting in higher chances of survival. These results suggest that intraspecific competition within these species could play a major role in seedling establishment rate. This factor needs to be taken in consideration when planning for seeding operation, to avoid overseeding and wastage of valuable and expensive seeds [54].

### Demographic processes

In field experiments, soil conditions at the time of germination and emergence (**Error! Reference source not found**.) were suitable for the germination of these temperate grass species. Differently to what was described by James et al. (2011), where the major bottleneck in seedling recruitment was detected at the emergence phase (when germinated seeds failed to push through the soil), in this experiment, the drop between germination and emergence was relatively small with probability of emergence from germinated seed ranging from 0.92 in *A. scabra* to 0.61 in *R. geniculatum* (**Fig 6**). This trend might be due to the favourable climatic and soil conditions during the year the study was conducted, with average night and daily temperature ranging between 10° C and 18° C, and maintained soil moisture content of 0.08-0.18m^3^/m^3^ (water potential range between −0.2 and −0.7 MPa) during the first month after sowing, when most of the emergence occurred. These conditions have not allowed for the detection of the stress reduction proprieties of SA that were originally hypothesised at the germination and emergence phase. However, the field data, combined with the controlled germination experiment with reduced water availability, suggest that SA might not affect seed performances at the establishment phase, as suggested by [34]. Further studies are needed to test this hypothesis under more severe stress conditions and on different species.

**Fig 4.**
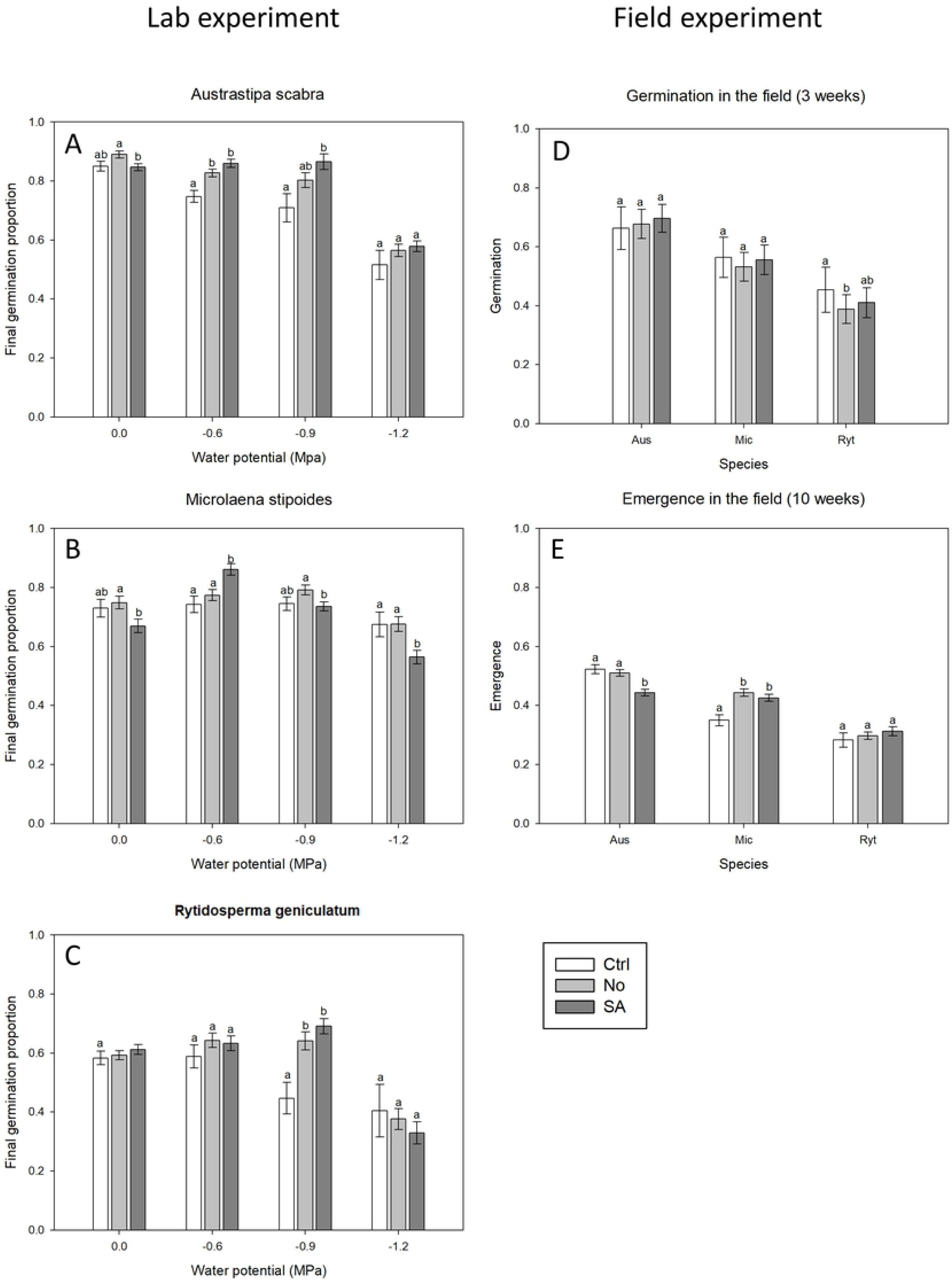
Salicylic Acid final germination and emergence. Final germination and emergence of untreated seeds (Ctrl), seed treated without salicylic acid (No) and seed treated with salicylic acid (SA). A, B and C shows the laboratory germination experiment in petri dishes at 20°c at different water potentials (X axis). D, E shows the germination and emergence results in the field experiment, 3 and 10 weeks after sowing respectively. The species are listed in the × axis (Aus = *Austrostipa scabra*, Mic = *Microlaena stipoides*, Ryt = *Rytidosperma geniculatum*). Results followed by the same letter for the Water potential (lab experiment) and species (Field experiment) are not statistically different at p < 0.05

**Fig 5.**
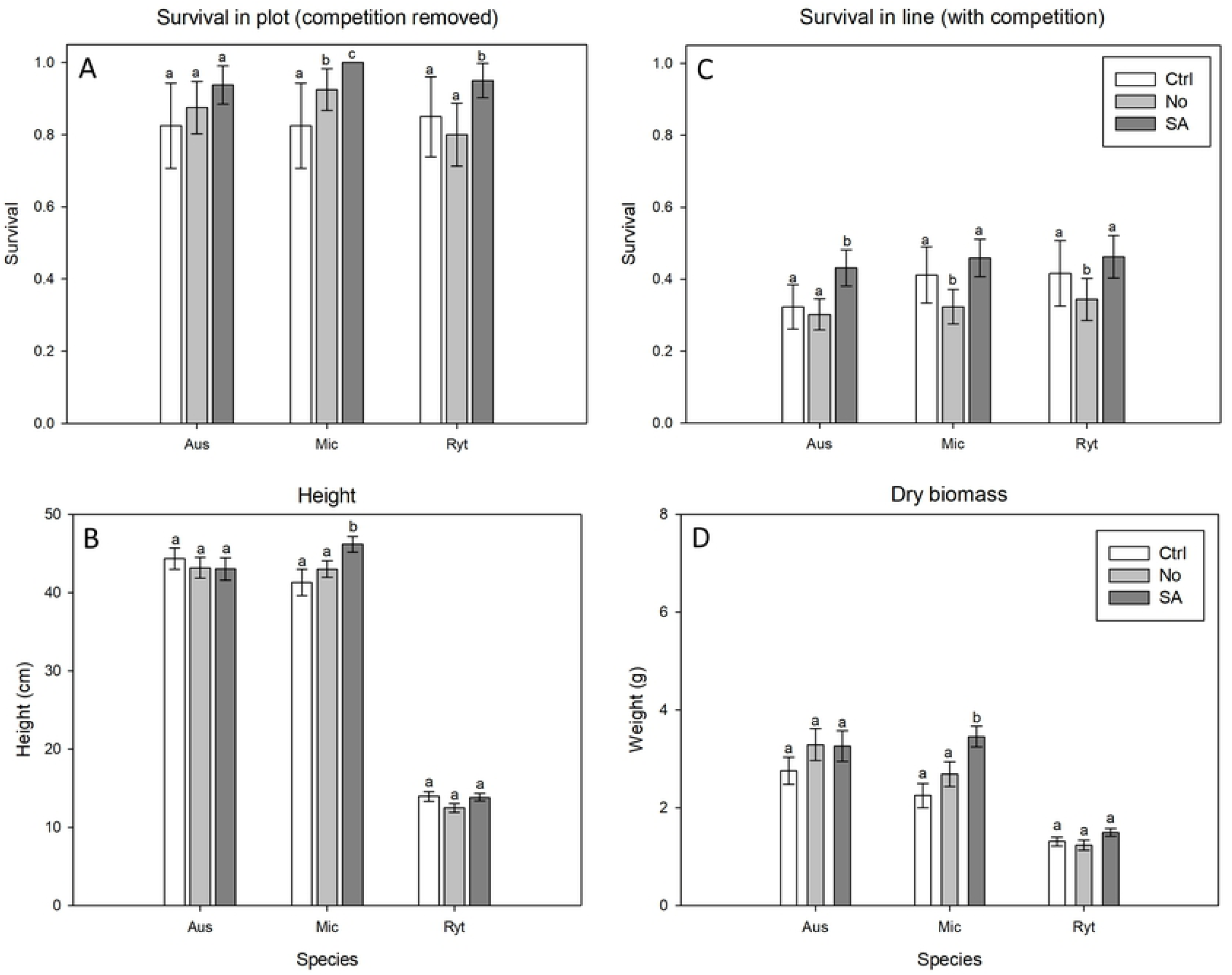
Survival and plan. Survival and plant growth comparison 40 weeks after sowing, between untreated seeds (Ctrl), seed treated without salicylic acid (No) and seed treated with salicylic acid (SA). (A) plant survival proportion in the plot experiment, where interspecific competition was limited, by removing excess seedlings and leaving 10 seedling per 0.25 m ^2^ plot. (C) Seeds sown on a 1 m line, without thinning. (C) Average height and (D) biomass of plant collected from the plot experiment. Results followed by the same letter are not statistically different at p < 0.05.

**Fig 6.**
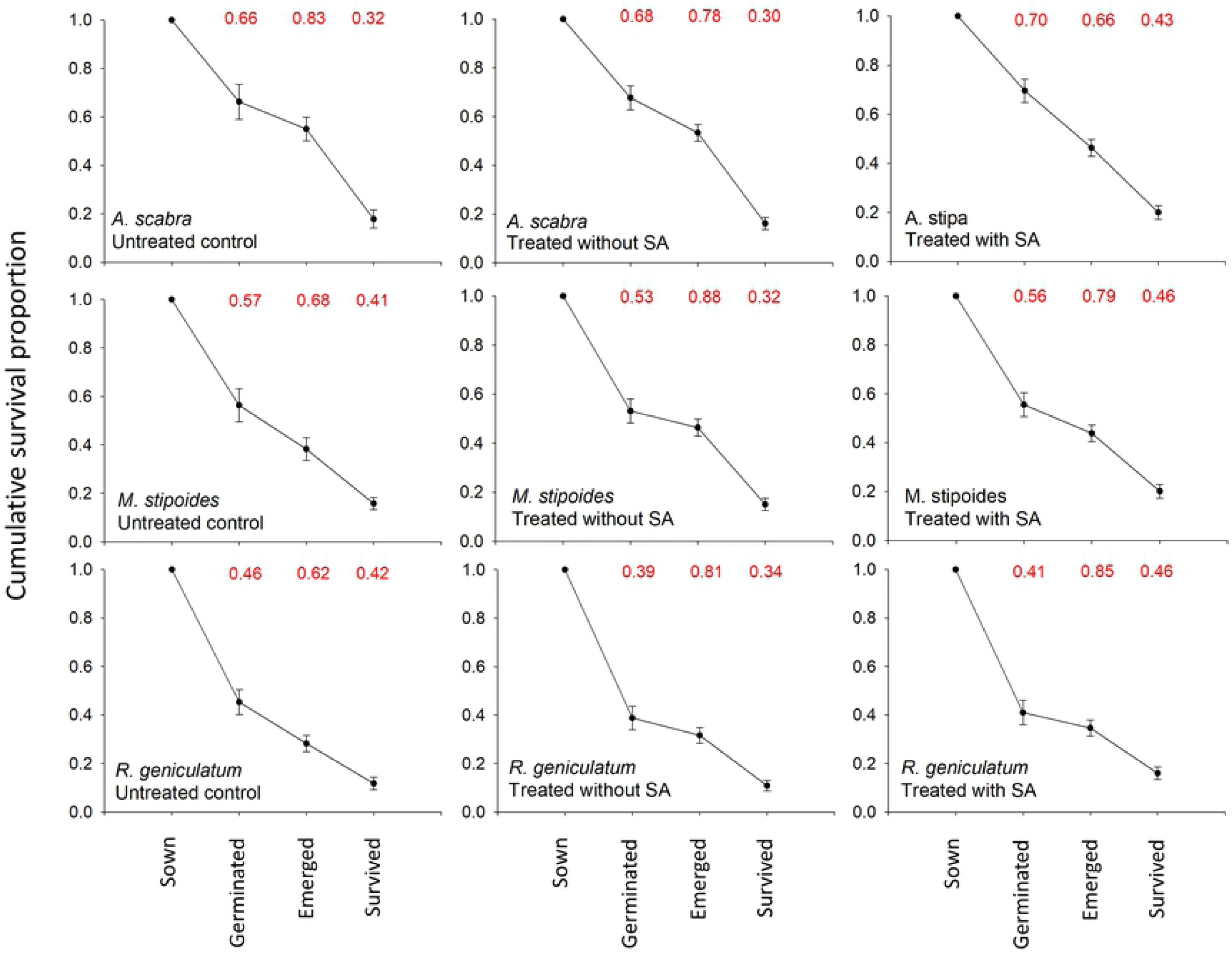
Cumulative survival proportion. Demographic process through various life stages for the three species tested without treatment, treated without SA and treated with SA. On the top of each graph, in red, are reported the probability of transitioning between life stages. This demographic data are based on the “in line’ experiment whereas seedling were not removed after emergence and intraspecific competition affected plant survival.

Significant effects of SA delivering stress resistance were instead detected on the survival of established plants over the summer when seedlings had to endure prolonged periods with little access to water. Total precipitation between November 2017 and February 2018, removing two major rainy events that happened over a short period (60 mm on December 20^th^ and 147 mm on January 18^th^) were less than 30 mm (Fig 2). The effects of the summer drought were evident on the experiment where seedlings were not removed, with the probability of plant survival from an emerged seedling being 0.32 for *A. scabra*, 0.41 for *M. stipoides* and 0.42 for *R. geniculatum*. In this case, SA treated seed survived significantly better than the seed treated without SA for the three species. When considering the cumulative survival from the number of seeds initially sown, SA treatment provides a significantly higher number of successful plant establishment events, even for *A. scabra*, when emergence of SA treated seed was lower than the seed treated without SA.

### SA effect on survival

In both line and plot experiments, SA treated seeds improved survival, supporting previous evidence that SA exogenous application may deliver drought stress resistance [43]. This improvement in survival might be due to a variety of factors, such as the effect of SA in mediating reactive oxygen species (ROS) and triggering defence-related processes [55], and its effect on productivity and growth [56]. In this study, just one of the three species tested (*M. stipoides*) showed a higher biomass production as a response to SA treatment. A previously published study reported that externally applied SA had increased root development [57], but root growth was not evaluated in this study. Nevertheless, as this study shows, the effects of exogenous SA delivery are still present months after its application. SA absorbed through the seed (imbibing), or through emerging radicle and roots (encrusting) could be converted in SA glucoside and transferred in the vacuole for storage [58]. SA glucoside could be mobilized and moved through the plant after been converted in methyl salicylate, and eventually turned back to SA when needed [27].

### Encrusting and imbibition

When SA delivery mechanisms of imbibing and encrusting were compared in terms of improving plant survival, a significant difference was rarely detected, suggesting that seed encrusting could be used to deliver SA and its stress resistance inducing proprieties. The advantage of using SA in the seed coating processes over imbibition lies in the capability of storing seed after treatment. Seed imbibition can trigger a seed priming effect that could improve germination speed and synchronicity in the short term [59], but, such imbibition could accelerate seed ageing processes, reducing seed shelf-life and storability [60]. Another advantage of seed coating over imbibition is that while it delivers SA stress resistance, it can also improve seed handling and sowability, along with a wide variety of active ingredients, such as protectants, micronutrients, germination promoters and microorganism [24]. Most of these coating treatments still need to be tested on native species for restoration, but their combined impact on seed germination, emergence, growth and plant establishment could improve the successful deployment of native seed onto degraded landscapes, ultimately allowing for a more cost-effective seed-based restoration.

## Acknowledgements

We acknowledge that the research undertaken and presented here was done on Wadjuk Noongar country and pay our respects to their elders, past, present and emerging.

This study is dedicated to the memory of the late Dr Tissa Senaratna who was instrumental in establishing salicylic acid as a key principle for improving plant growth and development and without whom this and many other studies would not have been possible.

S.P. was the recipient of a Curtin University International Postgraduate Research Scholarship. A.T.C is the recipient of the Research Fellowship in Restoration Ecology jointly funded by the EcoHealth Network, Gelganyem Limited, and Curtin University. Seed were donated by Native Seed Pty Ltd. This research was supported by the Australian Government through the Australian Research Council Industrial Transformation Training Centre for Mine Site Restoration (Project Number ICI150100041). The views expressed herein are those of the authors and are not necessarily those of the Australian Government or Australian Research Council.

## Supporting information

**S1_GerminationEmergenceAnalysisResults**.**pdf**. Final germination and T50 value of germination experiments of the three test species at full and reduced water potential and emergence in the field experiment. Statistics obtained with parameter comparison of DRM model comparing treatment, SA, and combination of treatment and SA against the untreated control.

